# Shedding Light on *H. pylori* Detection: A Fusion Protein Approach Unveiled through LIPS Method

**DOI:** 10.1101/2024.06.06.597801

**Authors:** Seyedeh Mahsa Farzanfar, Sedigheh Asad

## Abstract

The Luciferase Immunoprecipitation Systems (LIPS) method serves as a highly sensitive approach for quantitatively detecting antibodies to antigens, offering potential in identifying viral and bacterial infections. However, the substantial size of the luciferase-antigen fusion protein presents challenges in both production and folding. An alternative strategy employing epitopes rather than full length antigenic protein may circumvent issues associated with recombinant expression. *Helicobacter pylori*, a gram-negative bacterium, poses a risk of gastric cancer if untreated over time. This study focuses on the recombinant production of a fusion protein comprising in silico designed antigenic epitopes from the *H. pylori* urease protein and luciferase, aiming to reduce the fusion protein’s size and thus augment its expression in the *E. coli* system. By employing bioinformatic analysis, sequences encoding the antigenic regions were pinpointed and subsequently amplified via PCR. A luciferase-linker-epitope construct was devised and constructed accordingly. The *E. coli* Bl21 (DE3) strain was utilized to express the recombinant chimeric protein, which was subsequently purified to achieve a state of homogeneity. The molecular weight of the fusion protein was estimated to be 75 kilodalton. Verification of the chimeric protein’s proper folding and functionality was confirmed, as evidenced by a bioluminescence assay yielding an emission of 13.7 × 10^6^ (RLU/s). Furthermore, western blot analysis authenticated the fusion protein’s capability to bind specifically to *H. pylori* antibodies. These findings underscore the potential of the resultant protein as a promising candidate for *H. pylori* detection while also streamlining the recombinant production of LIPS fusion proteins.

**Key Points:**

- Epitope-driven protein design boosts *E. coli* expression for LIPS advancement.
- Improved *H. pylori* detection aids early gastric cancer identification.

## Introduction

Despite significant advances in preventing and treating microbial infections, their economic and social impact remains significant globally (Gerace et al. 2022; Deusenbery, Wang, and Shukla 2021). Early detection and efficient treatment of pathogens are imperative. Early diagnosis plays a vital role in enhancing patient outcomes, reducing healthcare costs, and mitigating the risk of antibiotic resistance (Vincent et al. 2015). The Luciferase Immunoprecipitation Systems (LIPS) method has emerged as a powerful technique for detecting microbial infections by capitalizing on the interaction between antibodies and bacterial or viral antigens (Burbelo et al. 2011). Utilizing luciferase-based assays, LIPS achieves highly sensitive and specific detection, exploiting the luminescent properties of luciferase to quantify pathogen presence with remarkable precision (Burbelo et al. 2010). Notably, LIPS surpasses conventional techniques like ELISA in sensitivity, enabling the detection of minimal antibody concentrations, and offers a broad dynamic range for assessing antibody levels across various titers (Burbelo et al. 2011). Its increasing adoption in diagnostic microbiology underscores its utility in rapidly identifying and characterizing microbial infections (Steffen et al. 2019).

LIPS technology has been used to diagnose various bacterial and viral infections such as *Strongyloides stercoralis, Onchocerca volvulus, Loa loa*, EBV, HIV, HTLV-1, HCV, *Wuchereria bancrofti*, African swine fever (ASF), KSHV, *Borrelia miyamotoi*, EBOV, *toxoplasmosis*, human norovirus (HuNoV), NPHV, and MERS (Burbelo, Lebovitz, and Notkins 2015; Bennuru et al. 2020; Aye et al. 2020; Ding et al. 2022; Tin et al. 2017). Since late 2019, the COVID-19 pandemic spurred scientists to utilize LIPS for studying humoral responses against SARS-CoV-2 (Hachim et al. 2020). These investigations have examined antibody emergence post-COVID-19 infection in various patient groups, identified autoantibodies in severe cases, tracked antibody evolution, screened antigenic protein responses, and assessed vaccine efficacy (Burbelo et al. 2020; Bastard et al. 2020; Dispinseri et al. 2021; Delmonte et al. 2021). These studies collectively advance understanding of SARS-CoV-2 infection dynamics, immune responses, and vaccine effectiveness (Atanackovic et al. 2022). Despite the numerous advantages outlined for the LIPS method, the production of the fusion protein combining antigen and luciferase frequently results in a large size, rendering its production unfeasible in prokaryotic expression system. Consequently, fusion protein production has predominantly occurred within the HEK 293 cell expression system or other cells such as Cos1 (Burbelo, Ji, and Iadarola 2023). To overcome this challenge, implementing epitopes instead of entire antigens might facilitate folding and expression of the fusion protein within *E. coli* cells, simplifying and streamlining the production process (Vita et al. 2019).

*Helicobacter pylori*, a gram-negative, spiral-shaped bacterium, stands as the leading cause behind peptic ulcers, gastric cancer, and various other ailments, including non-alcoholic fatty liver disease, iron deficiency anemia, and specific types of lymphoma (Matsunaga et al. 2018; Parkin et al. 2005). Despite often being asymptomatic, individuals exhibiting symptoms or harboring suspicions should undergo regular monitoring and diagnostic testing for *H. pylori* (Khan and Howden 2019; Vogelmann and Amieva 2007). Implementing the LIPS assay for detecting *H. pylori* could significantly enhance infection control and monitoring (Fig 1).

**Fig 1.**
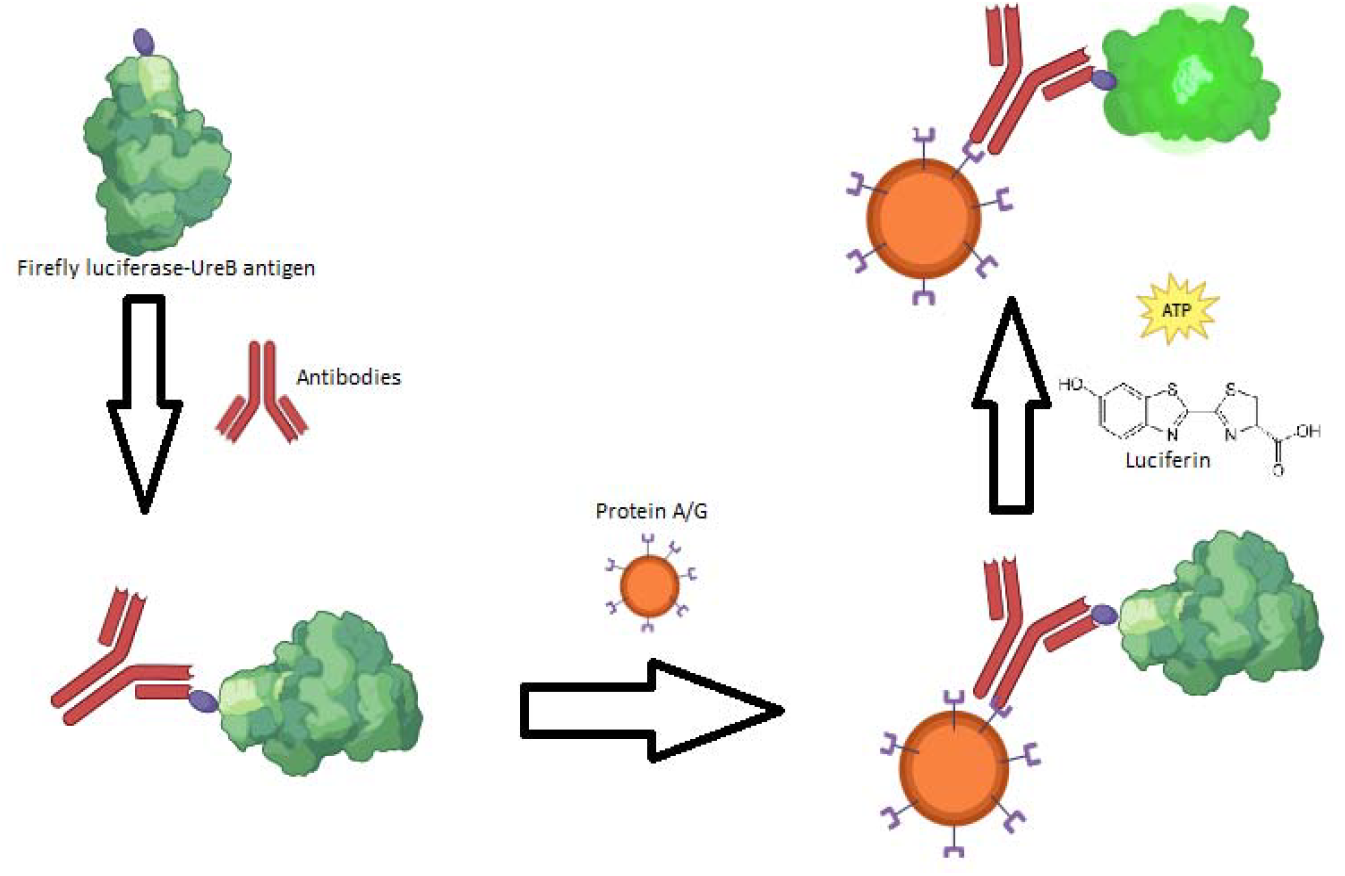
Schematic representation of the LIPS assay for the detection of H. pylori infection

Previous studies have developed a methodology to identify effective proteins serving as *H. pylori* antigens, with urease B subunit (UreB) emerging as a promising candidate (Ma et al. 2021). This antigen holds significant promise for both diagnostic and therapeutic purposes, given its highly conserved sequence across various *H. pylori* strains (Calado 2022). In this study, a luciferase fuged chimeric protein was designed and produced with linear epitopes of the UreB antigen in *E. coli*. The luciferase activity and binding to specific antibodies of the chimera were evaluated. The resulting fusion protein can be used for the development of LIPS as an efficient diagnostic tool for monitoring and controlling patients and individuals at high risk of developing this infection.

## Materials and Methods

### Chemicals, bacterial strains, and cell lines

The chemicals utilized in this investigation were all of analytical grade purity. Isopropyl-β-D-thiogalactopyranoside (IPTG) were purchased from Sigma Aldrich (USA). Primers were procured from Macrogen (South Korea). T4 DNA ligase and restriction endonucleases were acquired from Thermo Fischer (USA). Agarose, DNA ladder and PCR master mix were obtained from kawsarbiotech (Iran). The anti-UreB antibody was obtained from a rabbit immunized with the UreB antigen (Pasteur Institute, Iran). All remaining chemicals were procured from Merck (Germany), unless explicitly stated otherwise. *H. pylori* strains (NCTC 11637) were obtained from the cell bank of the Pasteur Institute (Iran).

### In silico studies

The process of designing the fusion protein comprised several stages. Initially, protein characteristics such as molecular weight, size, and theoretical isoelectric point (pI) of the urease beta subunit of *H*.*pylori* were estimated using UniProt and Expasy websites. Next, the NCBI BLAST algorithm was utilized to discern sequence similarities with other ureases. Following this, Three B-cell and MHCbinding epitopes, previously documented, were chosen (Ma et al. 2021; Vita et al. 2019). The identified antigenic epitopes were subsequently verified using Vaxijen v2.0. PyMOL was employed for visualization and identification of the structure and positioning of these epitopic segments on UreB 3D model (PDB: 6QSU). The coding sequence corresponding to the target epitopes was then utilized for construction of a genetic cassette (WP_000724308.1). In the design of the fusion protein, an engineered variant of firefly luciferase (FLuc) was employed. The epitopic segments were engineered to be linked to the C-terminus of luciferase via a Gly-Ser-Ser-Ser-Lys-Leu-Ser linker. The resulting chimeric protein’s three-dimensional structure was modelled using I-TASSER server (Zheng et al. 2021; Zhang, Freddolino, and Zhang 2017; Yang and Zhang 2015), and its physicochemical properties were forecasted using the Protparam server (Garg et al. 2016). Epitopes in the fusion protein 3D model were predicted using the ELLIPro server (Ponomarenko et al. 2008).

### Molecular Cloning

A luciferase-linker-epitope construct was devised for insertion into the pET21a expression vector, known for its robust T7 promoter and N-terminal hexahistidine tag. The linker sequence, GGTTCTTCTTCTAAGCTTAGC, was incorporated into the primers designed for both firefly luciferase (Fluc) and UreB epitope cloning. The forward and reverse primers for Fluc gene amplification (previously cloned in the pET26 vector) were GGCTATACATATGGGCAGCTCTCATCATCATC and GCTAAGCTTAGAAGAAGAACCCATTTTGGCAACC, with the corresponding NdeI and HindIII restriction sites marked. The PCR product obtained was purified, subjected to double digestion, and subsequently ligated into the pET21a vector through conventional methods before being utilized for transformation. Clones harboring the firefly luciferase gene were selected for further manipulation.

The genomic DNA of *H. pylori* was isolated using the phenol-chloroform method (Tan and Yiap 2009). A supposed DNA fragment that encodes the antigenic determinant of the ureB gene (450 bp) was PCR amplified using primers GGTTCTTCTTCTAAGCTTAGCTTAGCGGATCAAATTG and TTATCTCGAGTTAACCCACACGACCCATCG, which included HindIII and XhoI restriction sites (underlined). Following enzymatic digestion, the PCR product was inserted into the recombinant vector that contained the Fluc gene. Later on, through the process of restriction analysis and sequencing, clones that possessed the accurate Fluc-linker-ureB epitope construct were identified (Macrogen, Korea).

### Protein Expression and Purification

*E. coli* BL21 (DE3) cells, harboring the recombinant vector, were cultured in 150 ml of Luria-Bertani (LB) broth medium. The culture was initiated with a 1% inoculum derived from bacteria grown overnight, and 100 µg/ml ampicillin was added as a supplement. The culture was subsequently placed in an incubator set at 37°C and agitated at 160 rpm until it reached an optical density of 0.5-0.6. Varying IPTG concentrations (0.1, 0.3, 0.5, and 1 mM), induction temperatures (30 and 20°C), and induction times (3, 5, and 20 h) were investigated to enhance the efficiency of fusion protein expression and simultaneously inhibit protein aggregation. Ultimately, protein expression was initiated by the addition of 0.5 mM IPTG for a duration of 20 h at a temperature of 20 °C.

Following induction, the cellular sediment was collected via centrifugation at 9000 g and 4°C for 10 min and suspended again in 3 ml of chilled lysis buffer containing sodium phosphate buffer 50 mM, NaCl 300 mM, and 2 M urea (pH 7.6). To further disrupt the cells and release their contents, the reconstituted pellet was subjected to sonication for a period of 10 min, where sonication was performed in 10 s pulses followed by 5 s resting intervals using a SYCLON Ultra Sonic Cell sonicator. Upon completion of the cell lysis process, the resulting lysate was subjected to centrifugation at a speed of 9000 g (for 30 min at 4°C). subsequently the supernatant was applied onto a Ni-NTA agarose column that had been pre-equilibrated with the lysis buffer. To elute the fusion protein, a sodium phosphate buffer containing 100 mM imidazole was employed. The fractions that contained the desired fusion protein were pooled. Imidazole removal and protein concentration was achieved through an Amicon centrifugal filter unit. The process of purification was performed at a temperature of 4°C.

Protein expression and purification steps was investigated by SDS-PAGE (12% w/v acrylamide) using standard laboratory protocols (Sambrook and Russell 2001). Utilizing the Bradford method, the protein concentration was estimated by employing bovine serum albumin as the standard using a Perkin Elmer lambda 25 UV/VIS spectrophotometer (Bradford 1976).

### Fusion protein efficiency studies

#### Luciferase Activity Assay

To assess Fluc activity, we incubated 60 μL of the purified fusion protein solution (0.07 mg/ml) with 78 μM luciferin in a buffer comprising 0.66 mM Dithiothreitol (DTT), 25 mM Tris acetate, 11 mM Magnesium acetate, 1.1 mM bovine serum albumin (BSA), and 1.1 mM ethylenediaminetetraacetic acid (EDTA) for 20 s at 28°C. Subsequently, 0.1 mM ATP solution (pH 7.8) was added and the emitted light was measured by a Berthold luminometer (Germany).

#### Antibody binding assay

Western blot analysis was conducted to assess the binding of the chimeric protein to its specific antibody. The SDS-PAGE gel was loaded with 20 µl of protein samples, which were then subjected to electrophoresis. Following this, the samples were transferred to a PVDF (polyvinylidene difluoride) membrane using wet transfer method. The membrane was then blocked with 1% BSA in 0.5% Tween-PBS. Primary antibody, rabbit anti-UreB antibody, diluted 1:2000, was incubated with the blots and left at 4°C overnight. subsequently, the blots were exposed to a secondary antibody, specifically horseradish peroxidase conjugated goat anti-rabbit IgG which was diluted to a ratio of 1:15000. This treatment was carried out at room temperature for a duration of 1 h. Following four 5-min washes in TBST (Tris-buffered saline, 0.1% Tween 20), the membrane was prepared for visualization of protein bands using ECL (Enhanced Chemiluminescence) substrates.

## Results

Three B cell epitopes have been previously documented within the *H. pylori* urease protein, specifically situated on the enzyme’s beta subunit (see Table 1).

**Table 1.**
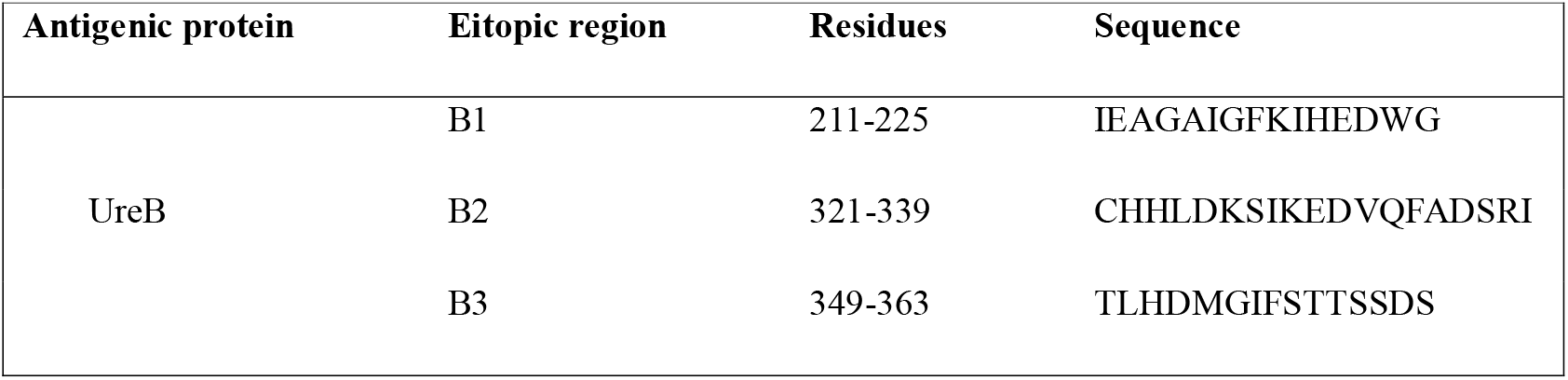
Antigenic regions of UreB gene that are detected by B-cell produced antibodies (Ma et al. 2021)

These epitopes exhibit a high degree of conservation across various ureases. The spatial location of these epitopes is illustrated in red within the enzyme’s 3D model (Fig 2).

**Fig 2.**
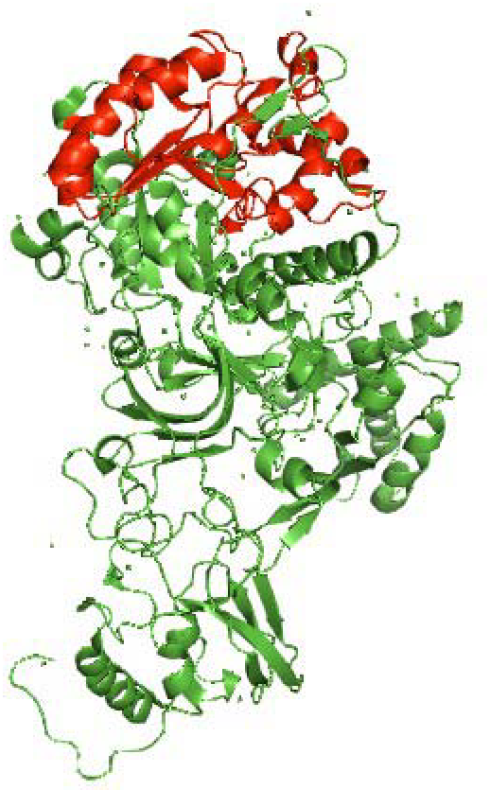
Antigenic Regions of UreB (WP_000724308.1 EC 3.5.1.5) Visualized in PyMOL Software. The selected antigenic regions are highlighted in red color for clarity and emphasis

The identified epitopes underwent validation using Vaxijen and ELLIPro, assessing both the primary and tertiary structure of the enzyme. Given that these three epitopes are linear in the urease primary structure, we opted to incorporate the protein segment spanning residues 206-365, encompassing all three epitopes. For the design of the fusion protein, we selected an engineered variant of firefly luciferase accessible in our laboratory. To connect the two components, a series of Gly-Ser residues were engineered as linker. In silico analysis estimated the molecular mass of the FLuc-linker-UreB epitopes fusion protein (733 aa) to be approximately 75 kDa. The protein’s theoretical isoelectric point (pI) was predicted to be 5.94. Furthermore, the predicted 3D model indicated that the protein exists as a monomer with a predominantly hydrophilic surface. Subsequently, the presence of B-cell epitopes was reconfirmed within the chimeric protein’s 3D model (Fig 3).

**Fig 3.**
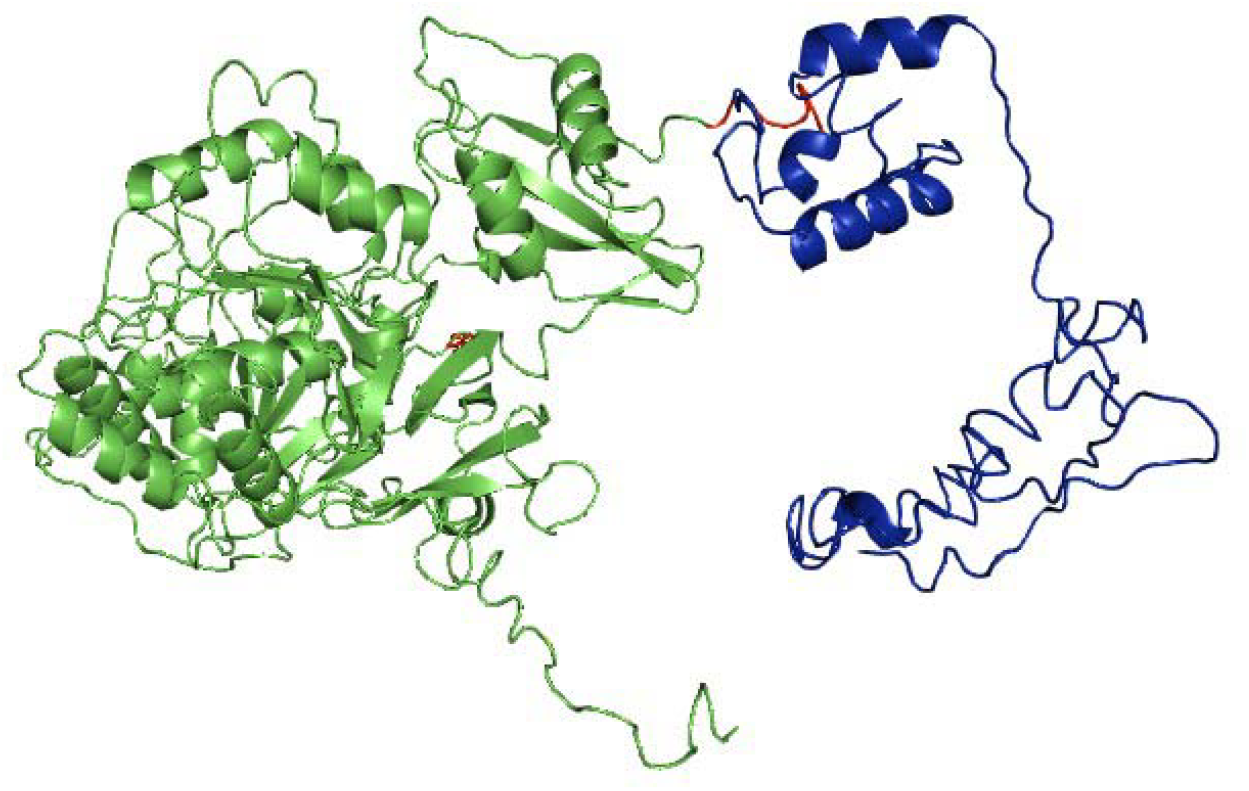
3D Structure of Fusion Protein: the three-dimensional structure of the FLuc-ureB epitope fugion protein, generated using the I-TASSER server. The model provides insights into the spatial arrangement and conformation of the fusion parts

A 1710 base pair fragment containing the FLuc coding sequence was successfully amplified from the pET26+luciferase plasmid via PCR. This fragment was then cloned into the pET21a vector, where it was positioned downstream of the hexa histidine coding sequence. The resulting recombinant vector, now containing FLuc, was used for the subsequent cloning step. Simultaneously, the coding sequence corresponding to the ureB epitopes and flanking regions, a 526 base pair fragment, was PCR amplified from the *H. pylori* genome and integrated into the recombinant plasmid. This process yielded a chimeric sequence 2202 base pairs in length (Fig 4 and 5). The sequencing analysis revealed that both fragments aligned precisely with the intended designs and importantly, a complete open reading frame was created.

**Fig 4.**
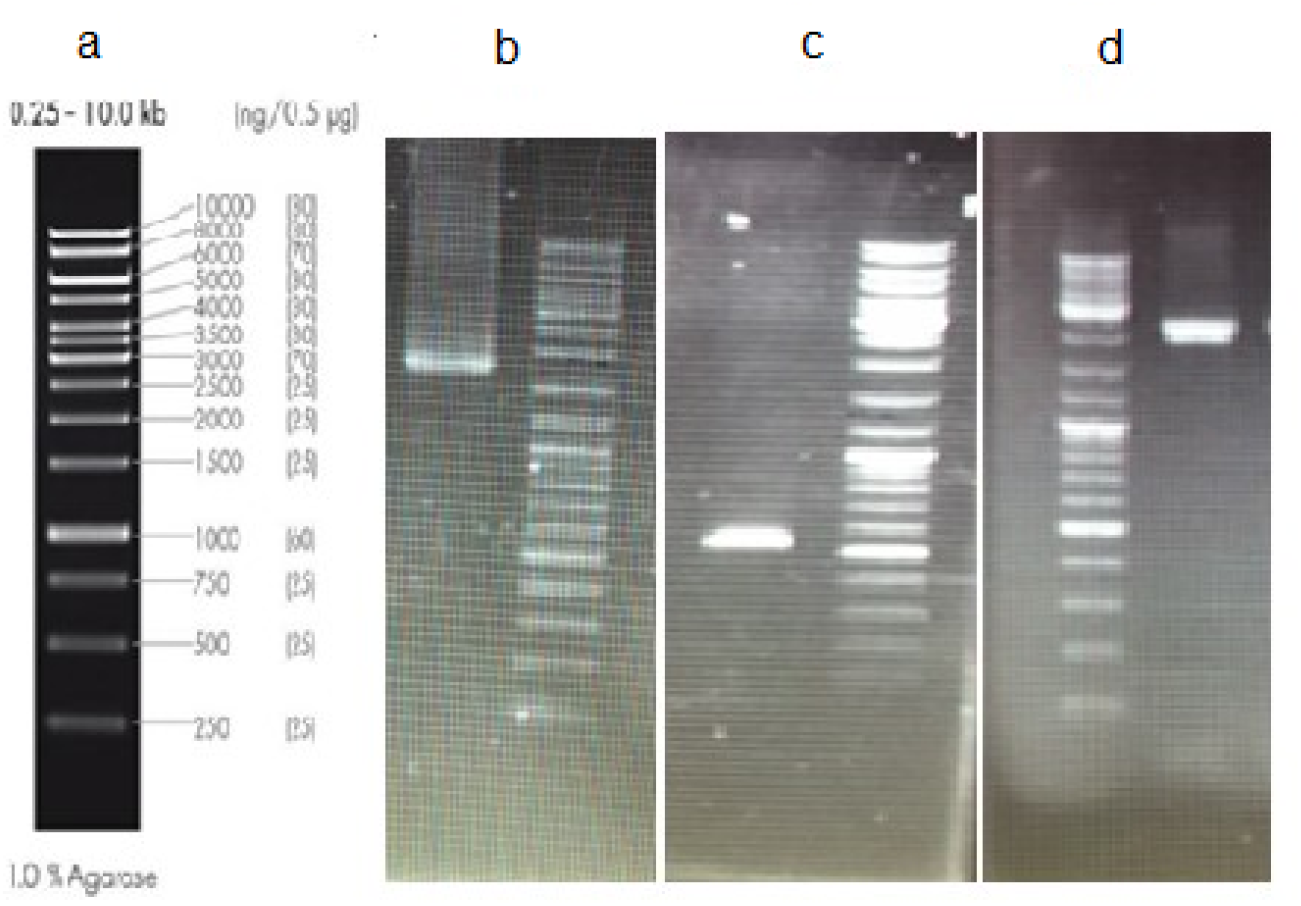
Agarose gel electrophoresis of the PCR-amplified fragment with a ladder map. a: DNA ladder, b: PCR product of the luciferase (1710 bp) c: PCR product of the UreB epitopes (526 bp) d: PCR product of the FLuc-linker-UreB epitope cassette (2202 bp)

**Fig 5.**
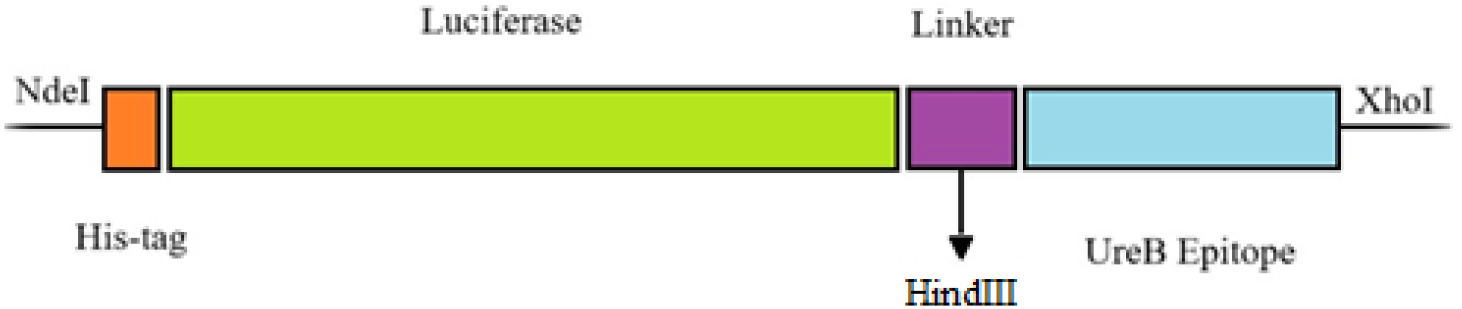
Recombinant chimeric genetic cassette, a visual representation of the arrangement and organization of the constituent genes within the chimeric cassette

Expression of the chimeric protein was conducted in *E. coli* BL21(DE3) utilizing IPTG (0.5 mM) and a 5-h induction period at 30°C. Analysis via SDS-PAGE shown a high level of expression of the recombinant protein; however, it predominantly appeared as inclusion body. To overcome this challenge, optimization experiments were conducted with various concentrations of IPTG, induction temperature and duration, as described in the methods section. Through these optimizations, lowering the IPTG concentration to 0.5 mM and prolonging the induction period to 20 h at a reduced temperature of 20°C significantly enhanced the soluble production of the desired fusion protein. Subsequently, the fusion protein was purified to homogeneity using immobilized metal affinity chromatography on a nickel-agarose column. The chimeric protein migrated on SDS-PAGE with a molecular weight corresponding to its predicted size of 75 kDa (His-tagged protein) (Fig 6).

**Fig 6.**
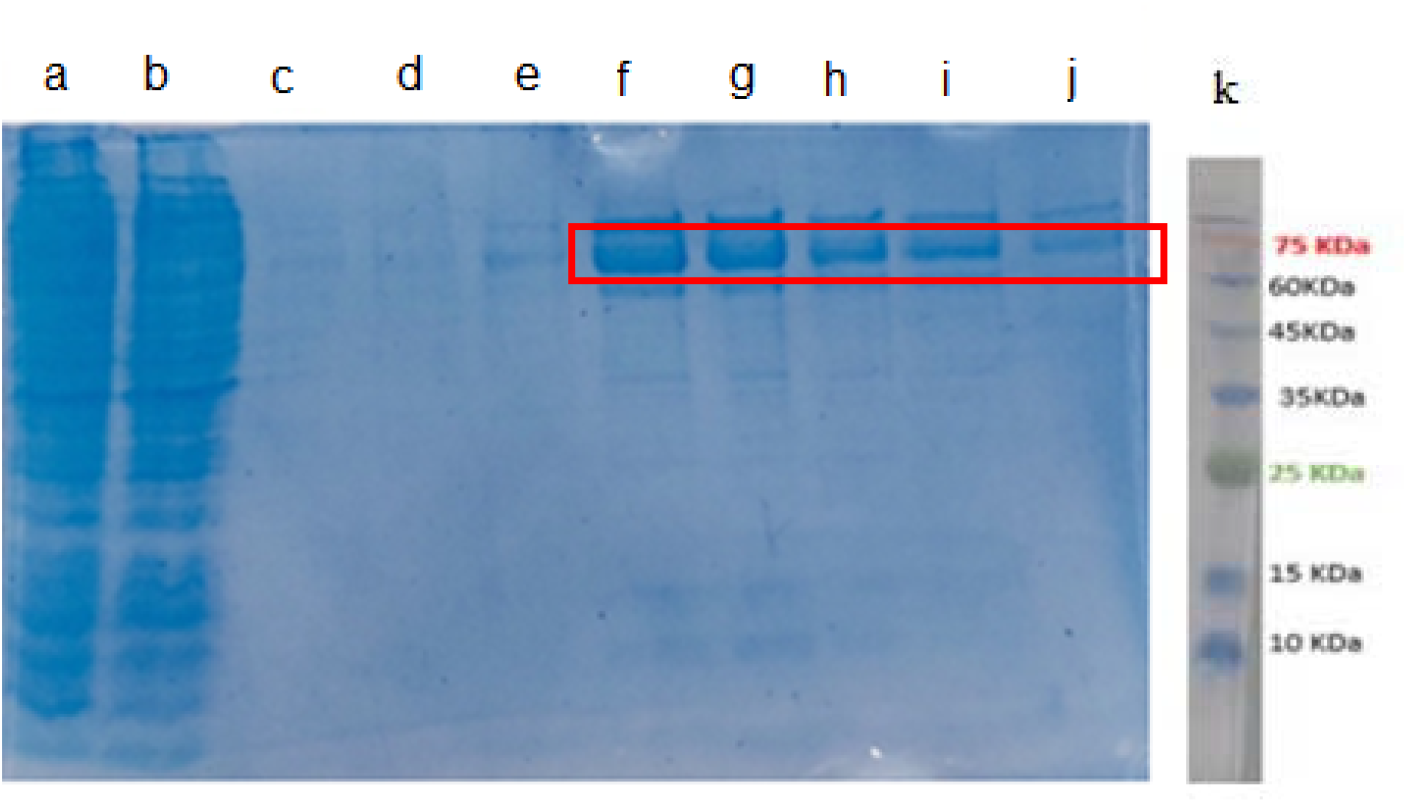
Purification of protein with histidine tail with the help of a nickel column, this figure in order from left to right: a) Total proteins after induction, b) Flowthrough, c) Washed proteins, d-j) Elution fractions, k) Protein ladder

The chimeric protein generated a luminescent signal upon incubation in the reaction buffer containing luciferin and ATP. Utilizing a luminometer, the emitted signal was quantified and yielded 13664710 Relative Luminescence Units per second (RLU/s). Western blotting analysis was conducted to evaluate the interaction between the chimeric protein and UreB specific antibody, with full-length UreB antigen serving as a positive control for comparison. Additionally, the binding of the chimeric protein to serum samples obtained from both healthy individuals and those infected was examined. As depicted in Fig 7, the chimeric protein exhibited successful binding to the specific antibody, akin to the full-length UreB antigen. The disparity in size observed is attributable to the larger molecular weight of the chimeric protein relative to the UreB antigen (Karimi and Mohammadi 2001). Notably, in the serum sample derived from the infected individual, a band at approximately 75 kDa was identified, indicating the chimeric protein’s capability to distinguish infected individuals. Conversely, in the serum sample obtained from the healthy individual, no corresponding band was noted but a smaller band, approximately at 60 kDa, was observed.

**Fig 7.**
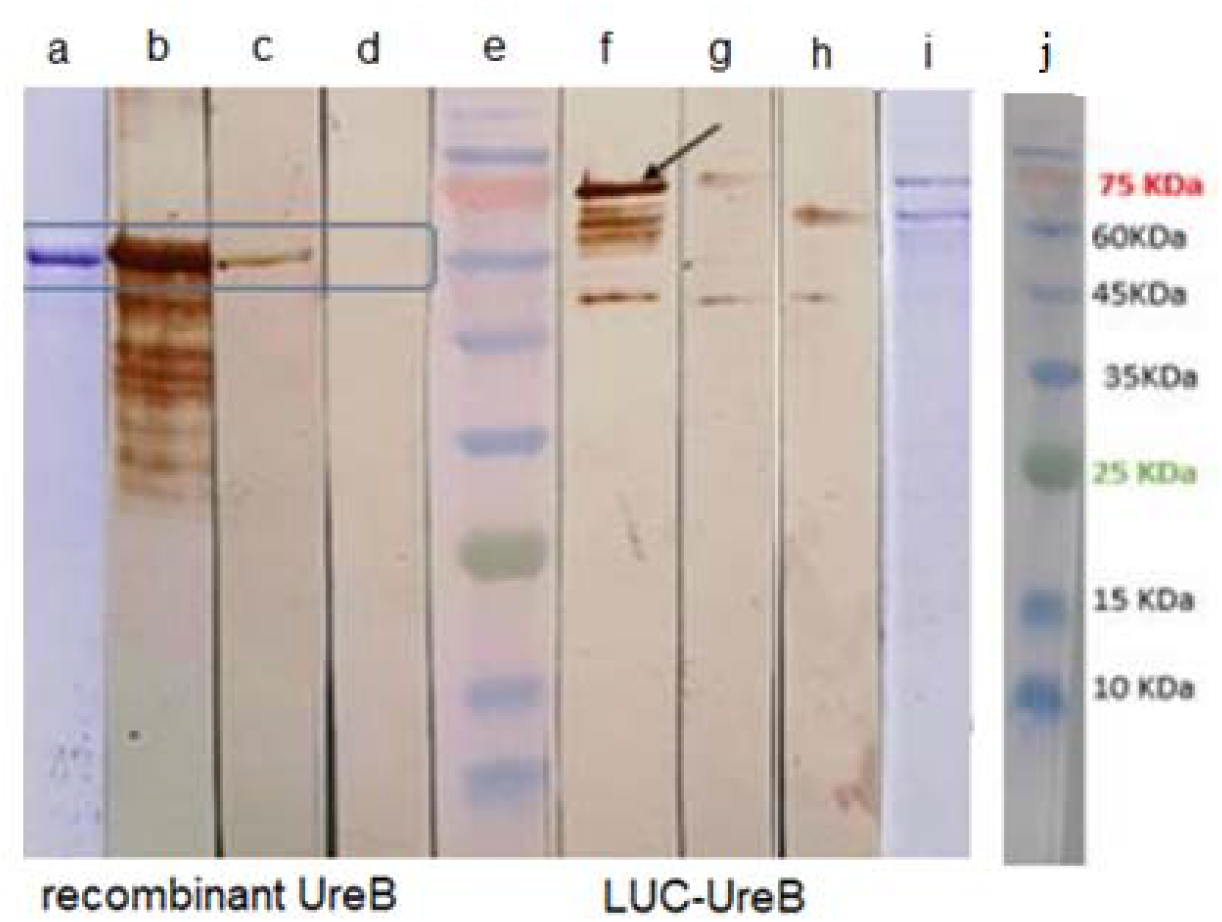
Western blot Analysis of UreB Protein and Recombinant Chimeric Protein. a: SDS-PAGE gel of UreB protein, b: Western blot of UreB protein using the specific antibody, c: Western blot of UreB protein using blood serum from a sick individual, d: Western blot of UreB protein using blood serum from a healthy individual, e: Molecular weight ladder, f: Western blot of our recombinant fusion protein using the specific antibody, g: Western blot of the recombinant fusion protein using blood serum from a sick individual, h: Western blot of the recombinant fusion protein using blood serum from a healthy individual, i: SDS-PAGE gel of the affinity IMAC purified fusion protein, j: protein ladder map

## Discussion

LIPS technology addresses the urgent need for presice antibody measurement in autoimmune and infectious diseases. Its pioneering use of luciferase fusion proteins has led to significant advancements, including its recent application by the FDA in evaluating the RSV vaccine’s efficacy (Kumari et al. 2014). This method boasts several advantages, including high specificity and rapid assay development (Burbelo, Ji, and Iadarola 2023). *H. pylori* infection presents significant health challenges, contributing to various conditions like gastritis, gastric ulcer, and iron deficiency anemia (Parkin et al. 2005; Du and Atherton 2006). While eradicating this infection offers benefits such as healing ulcers and preventing certain cancers, current diagnostic methods, including biopsy and serological tests, either involve invasiveness or lack accuracy (Rimbara, Sasatsu, and Graham 2013). Developing a LIPS system tailored for *H. pylori* detection promises a simple, sensitive, and non-invasive diagnostic approach.

The current investigation aimed to develop a Luciferase Immunoprecipitation System (LIPS) for detecting *H. pylori*, necessitating the selection of a conserved and immunogenic antigen. Given the prevalence of urease antigen in clinical trial vaccines, particularly exemplified by Zeng et al.’s 2015 urease-based pylori vaccine, which showed promise in phase III trials, the UreB antigen was chosen for this purpose (Angelakopoulos and Hohmann 2000; Sougioultzis et al. 2002; Aebischer et al. 2008; Zeng et al. 2015). However, producing the desired luciferase-UreB fusion protein in the *E. coli* expression system is challenging due to its size and folding constraints. Consequently, alternative cell lines such as HEK 293 were employed in these studies, albeit at a higher cost and with greater difficulty. Nonetheless, this study circumvented this issue by utilizing epitopes of the UreB antigen instead of whole protein.

During the in silico design process, the study initially opted for the protein segment encompassing all three epitopes identified in the work by Ma, J., J. Qiu et al (Ma et al. 2021). A linker sequence was then strategically designed between the UreB epitopic fragment and FLuc sequences to ensure proper folding and activity of the fusion protein. The epitope coding region, extracted from *H. pylori*, was successfully cloned alongside luciferase in pET-21. Sequencing confirmed the match between the obtained nucleotide sequence and the designed chimeric construct. Optimization of expression conditions enhanced soluble fugion protein production. Purification via Ni2+ affinity chromatography yielded an approximately homogenic protein with a molecular weight of 75 kDa estimated by SDS-PAGE.

In the scenario involving fusion proteins, the precise folding of every fragment holds paramount importance. Therefore, to assure the accurate folding of luciferase and epitopes, it was imperative to evaluate luciferase enzymatic activity and epitope’s antibody binding efficiency. Assessment of enzyme activitiy revealed preserved luciferase structure post-fusion, with the fusion protein exhibiting robust activity. Additionally, the antigenic region demonstrated specific antibody binding ability and elicited a positive response in infected individual serum samples in western blot analysis, affirming its functionality. The presence of an smaller band in the case of healthy human serum, is likely attributed to impurities present in the protein sample. Nonetheless, the absence of the chimeric protein band indicates the potential utility of the chimeric protein for diagnostic applications. Overall, these findings suggest that the fusion protein combining luciferase with the ureB antigenic region shows promise as a viable candidate for the advancement of LIPS as an efficient primary diagnostic approach. Nonetheless, additional experiments are warranted to validate its efficacy further.

This study represents the fisrt-ever documentation of a fusion protein engineered explicitly for the development of a Luciferase Immunoprecipitation System (LIPS) tailored for detecting *H. pylori*. This fusion protein introduces a novel approach for *H. pylori* diagnosis through integration into the LIPS methodology. It is noteworthy that no prior LIPS luciferase fusion protein incorporating epitopic regions rather than entire antigenic proteins has been reported, necessitating the production of fusion proteins within eukaryotic systems. In a 2023 investigation, HEK 293T cells were utilized for fusion protein production to evaluate Deltacoronavirus porcine (PDCoV), a coronavirus with potential cross-species infectivity (Boley, Lemon, and Kenney 2023). Similarly, in another study, HEK293FT cells were employed for producing recombinant HA proteins tagged with NLuc reporters for influenza virus HA detection (Honda et al. 2021). Furthermore, a LIPS assay for detecting IgG antibodies against hRSV’s G-glycoprotein was established using COS-1 cells for fusion protein production (Crim et al. 2019). Additionally, Christine M. Tin et al. developed a LIPS assay for defining antigenic specificity and antibody titers against HuNoV, utilizing COS1 cells for protein expression and transfection (Tin et al. 2017). In a 2022 study, HEK293T cells were utilized to develop a novel LIPS methodology based on the pREN2 plasmid containing the SFTSV NP coding sequence tagged with Rluc, investigating antibody responses to SFTSV (Chen et al. 2022). However, in this current study, the fusion protein was expressed in *E. coli* cells, resulting in a more cost-effective and streamlined process. Considering the widespread occurrence of *H. pylori* infection and the necessity for continuous monitoring of treatment efficacy and infection status, the creation of this rapid diagnostic technique holds immense potential to influence disease management and control significantly.

## Statements and Declarations

### Author contribution

S.M.Farzanfar: Methodology, Software, Validation, Formal analysis, Visualization, Writing original draft

S. Asad: Conceptualization, Supervision, Writing - Review & Editing, Project administration, Funding acquisition

### Ethics approval

This article does not include any research involving human participants or animals conducted by any of the authors.

### Data availability

The authors declare that they are aware of the availability of data in the main manuscript file.

